# Large-scale structural rearrangements unleash indiscriminate nuclease activity of CRISPR-Cas12a2

**DOI:** 10.1101/2022.06.13.495754

**Authors:** Jack P. K. Bravo, Thom Hallmark, Bronson Naegle, Chase L. Beisel, Ryan N. Jackson, David W. Taylor

## Abstract

Cas12a2 is a CRISPR-associated nuclease that performs RNA-guided degradation of non-specific single-stranded (ss)RNA, ssDNA and double-stranded (ds)DNA upon recognition of a complementary RNA target, culminating in abortive infection (Dmytrenko 2022). Here, we report structures of Cas12a2 in binary, ternary, and quaternary complexes to reveal a complete activation pathway. Our structures reveal that Cas12a2 is autoinhibited until binding a cognate RNA target, which exposes the RuvC active site within a large, positively charged cleft. Double-stranded DNA substrates are captured through duplex distortion and local melting, stabilized by pairs of ‘aromatic clamp’ residues that are crucial for dsDNA degradation and in *vivo* immune system function. Our work provides a structural basis for this unprecedented mechanism of abortive infection to achieve population-level immunity, which can be leveraged to create rational mutants that degrade a spectrum of collateral substrates.

## Introduction

Prokaryotic adaptive immunity typically utilizes CRISPR-Cas systems to target and degrade mobile genetic elements (MGE), including phage, transposons and plasmids^1,2^. However, it was recently discovered that Cas12a2 from *Sulfuricurvum* sp. PC08-66 instead relies on abortive infection (Abi) – that is, dormancy in response to the presence of an invader – to achieve population-level immunity, preventing the replication and transmission of MGEs (Dmytrenko 2022).

While Cas12a2 often co-occurs with Cas12a systems in bacteria and can utilize Cas12a crRNA, Cas12a2 recognizes an RNA target strand with a suitable protospacer-flanking sequence (PFS, 5’-GAAAG-3’) rather than the double-stranded (ds)DNA target of Cas12a (Dmytrenko 2022). Furthermore, Cas12a2 is immune to the effects of many anti-CRISPR (Acr) proteins that target Cas12a, and aside from a conserved RuvC nuclease domain and pre-crRNA processing region, Cas12a and Cas12a2 sequences bear little resemblance to one another (∼10-20%). Notably, Cas12a2 lacks a Nuc domain (involved in DNA target strand loading), but instead contains a zinc-ribbon and Cas12a2 contains a unique insertion domain in place of the Cas12a bridge helix.

Unlike many of the recently-characterized Abi systems^3–11^, Cas12a2 does not rely on the production of secondary messengers to achieve anti-phage immunity. Instead, Cas12a2 activation induces robust, nonspecific cleavage of single-stranded (ss)RNA, ssDNA, and dsDNA (Dmytrenko 2022) in *trans*. This mechanism is unique to Cas12a2, although the molecular basis for collateral nucleic acid degradation is unknown.

To understand the unique mechanisms of activation, substrate capture and indiscriminate nuclease activity underlying Cas12a2 function, we performed biochemical, structural, and in *vivo* analyses, including determining cryo-electron microscopy (cryo-EM) structures of autoinhibited Cas12a2-crRNA (binary complex) associated with an RNA target (ternary complex) and bound to both an RNA target and a double-stranded DNA collateral substrate mimetic (quaternary complex).

## Results

### Structure of Cas12a2 binary complex

To gain structural insights into Cas12a2 function, we first purified a binary complex consisting of catalytically active Cas12a2 and a mature crRNA and determined the structure using cryo-electron microscopy (cryo-EM) to a global resolution of 3.2 Å (**Extended Data Fig. 1**). The quality of the map allowed *de novo* modelling of the majority of Cas12a2, aside from the flexible PFS-interacting (PI) and zinc ribbon (ZR) domains.

Cas12a2 adopts a bi-lobed architecture (**Fig. 1**), with a recognition (Recognition, REC) lobe comprising the REC1 and REC2 domains and the nuclease (Nuclease, NUC) lobe consisting of the PI domain, wedge (WED), RuvC nuclease, ZR and the Cas12a2-specific insertion (“Insertion”). The overall architecture of the binary complex resembles an oyster, as opposed to the triangular “sea conch” shape of Cas12a^14^.

**Fig. 1.**
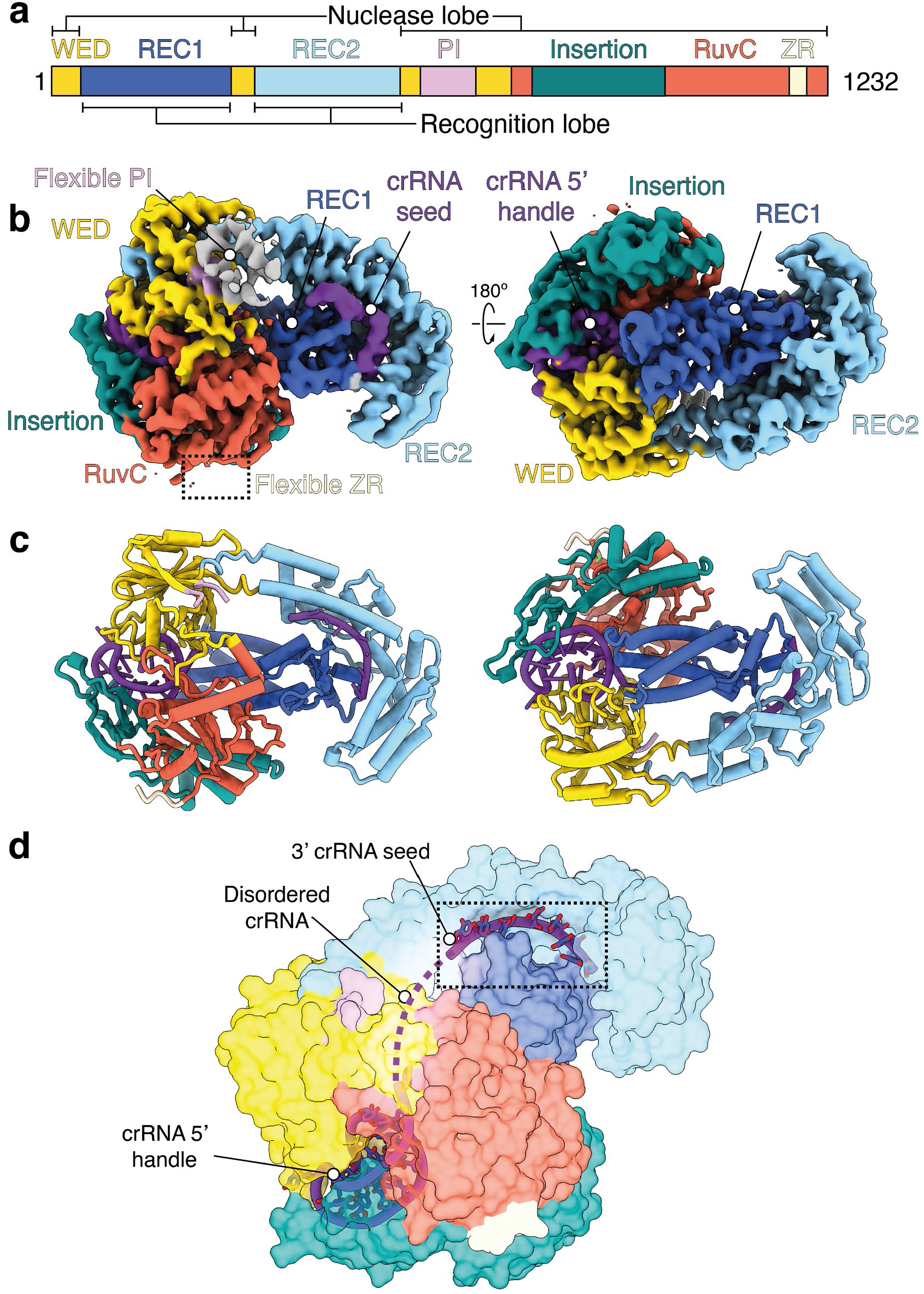
Cas12a2 binary complex resembles an oyster and orders the crRNA. **a**, Domain organization of Cas12a2. WED, Wedge domain; PI, PFS Interacting domain, ZR, Zinc Ribbon domain. **b**, Cryo-EM structure of Cas12a2 binary complex colored by structural domain as in (**a**). **c**, Atomic model of Cas12a2 binary complex. **d**, Putative 3’ seed region of crRNA. Seven bases from the 3’ end are ordered, and bases are solvent-exposed, likely acting as a seed for target RNA binding.

While Cas12a and Cas12a2 share low (10-20%) sequence similarity, comparison of the WED and RuvC domains shows a high degree of structural similarity (RMSD 1.073Å across 120 equivalent residues with FnCas12a (PDB 5NG6)) (**Extended Data Fig. 2**). Furthermore, the crRNA 5’ stem loop is in an identical configuration in both complexes, and a loop containing basic residues is similarly positioned to catalyze pre-crRNA maturation. The common “chassis” formed by these domains provides a structural scaffold that enables the same crRNAs to prime and guide either complex to the same target sequences for their different functions.

Despite the similar domain organization to Cas12a within the WED and RuvC domains, Cas12a2 has a unique α-helical REC lobe, with no known structural homologs. The differences in the structural organization of the REC lobe likely allow Cas12a2 to escape targeting by many anti-CRISPR (Acr) proteins that can efficiently shut down Cas12a (**Extended Data Fig. 3)**. Interestingly, 7-nts of pre-ordered crRNA sit at the interface between REC1 and REC2 in conformation where bases are solvent exposed and primed for target recognition (**Fig. 1d**). This region is towards the 3’ end of the crRNA, and the intervening sequence between the 5’ crRNA stem-loop and this pre-ordered guide is disordered in our structure, likely due to flexibility. This is in stark contrast to Cas12a, which has a well-described pre-ordered crRNA immediately flanking the 5’ stem-loop. In Cas12a, this region is highly sensitive to mismatches since it initiates R-loop formation upon PAM recognition and is considered a seed region^15^. In contrast, Cas12a2 is insensitive to single mismatches within the entirety of the crRNA but has reduced *in vivo* activity when truncated on 3’ end. Our structure suggests that Cas12a2 crRNA – target strand (TS) duplex formation may initiate and propagate from the 3’ end of the crRNA, enabling Cas12a2 to target phage that have escaped surveillance by Cas12a through mutagenesis of the PAM or 5’-seed regions.

### RNA target binding activates Cas12a2

To understand how RNA target recognition is distinct from other Cas12 family nucleases, we next investigated how RNA target binding activates Cas12a2. We determined a 2.9 Å-resolution cryo-EM structure of a ternary complex consisting of Cas12a2, crRNA, and a target ssRNA containing a non-self protospacer-flanking sequence (PFS, 5’-GAAAG-3’) (**Fig. 2**).

**Fig. 2.**
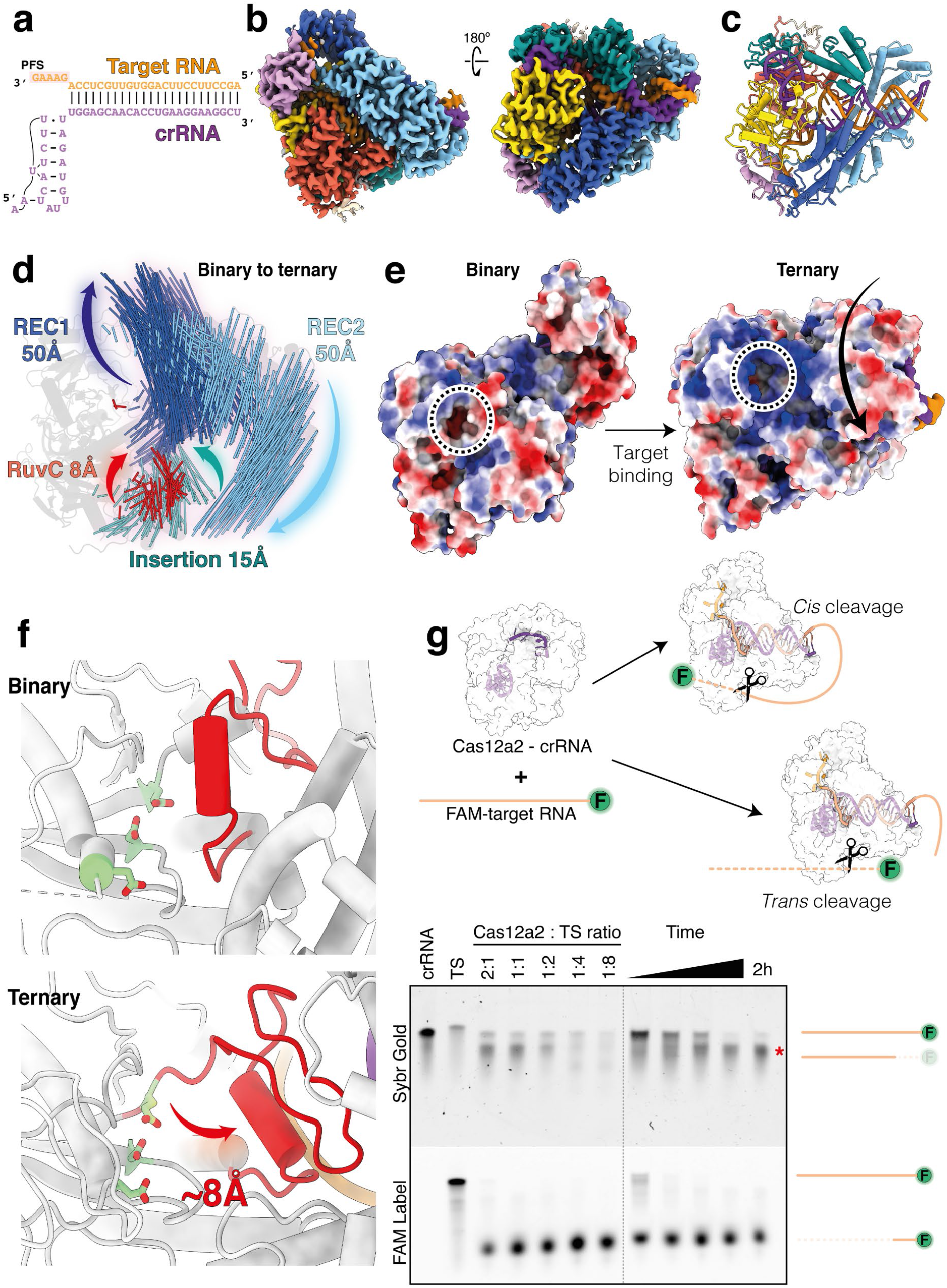
Target binding leads to large-scale structural arrangements for activation. **a**, Schematic of crRNA:TS duplex. Protospacer-flanking sequence (PFS) on target RNA is highlighted. **b**, Cryo-EM structure of Cas12a2 ternary complex. **c**, Atomic model of Cas12a2 ternary complex. **d**, Motion vector map showing conformational changes of Cas12a2 induced upon ternary complex formation. Binary complex model shown as grey cartoon. Conformational changes induced by target RNA binding are also shown in **Supplementary Movie 1. e**, Surface electrostatic potential of binary and ternary complex, showing how active site (dashed circle3) becomes exposed upon ternary complex formation. This is accompanied by the formation of a large positively-charged groove adjacent to the activate site. **f**, Displacement of RuvC gating helix (red) by ∼8Å upon ternary complex formation exposes active site residues (light green). **g**, Target protection revealed by Sybr staining, showing only peripheral cleavage of target RNA when binary complex is in molar excess, and total degradation when target is in excess. The observed protection persisted for 2H with excess binary complex.

The 22-bp A-form crRNA – target RNA duplex runs through the center of the complex. At the 5’ end of the crRNA guide, the duplex splits with the crRNA 5’ stem loop wedged between RuvC and WED domains, while the 3’ PFS end of the target RNA is gripped by the PI domain, which has now become ordered (**Fig. 2b**). Each of the 5-nts of the PFS make specific base contacts with residues within the PI domain, including hydrogen bonding and π-π stacking (**Extended Data Fig. 4**). These contacts stabilize the otherwise flexible PI domain, allowing Cas12a2 to distinguish self (i.e., complementary to the crRNA 5’ handle) from non-self target RNA based on the PFS. Removal of the PI domain had no effect on the overall structure of Cas12a2, but prevented activation of nuclease activity (**Extended Data Fig. 4**). Notably, this is a completely distinct mechanism of self-vs-non-self discrimination from Cas13 and several type III effector complexes, where nuclease activity is inhibited by additional complementarity with the 5’ crRNA tag region of the crRNA^16,18,19^. In contrast, Cas12a2 is exclusively activated upon recognition of an appropriate PFS sequence.

Cas12a2 is unable to degrade nucleic acids in the absence of a suitable target RNA (**Extended Data Fig. 4**). Superposition of the binary and ternary complexes reveals substantial conformational changes localized to the REC1 and REC2 domains, while the NUC lobe remains predominantly static (**Fig. 2d**). REC1 and REC2 are both displaced by up to ∼50 Å and move in different directions, creating a central channel, which accommodates the crRNA:TS duplex. The Insertion domain moves by up to ∼15 Å, but these changes are exclusively localized to the C-terminal half of the domain (residues 938-1030). The N-terminal half (residues 870-937), which makes numerous contacts with the crRNA 5’ stem-loop, remains static. Based on this observation, we propose that the Insertion domain acts as a transducer, allowing allosteric communication between the REC and NUC lobes in response to target RNA binding (**Extended Data Fig. 5**).

Inspection of the RuvC active site in the autoinhibited binary complex reveals that the catalytic triad (D848, E1063, D1213) is buried within a solvent-excluded pocket. Strikingly, the conformational changes that accompany target RNA binding create a 25 Å-wide positively-charged groove that exposes the active site (**Fig. 2e**). This groove is of sufficient size to accommodate both ss-and ds-nucleic acids. While other Cas12 proteins undergo conformational changes upon crRNA hybridization (up to ∼25Å in for Cas12a ^20^ and Cas12j ^21^, but more typically up to ∼10 Å^22^). The conformational rearrangements we observe for Cas12a2 are considerably larger, highlighting the distinct activation mechanism of Cas12a2 (**Supplemental movie 1**).

Access to the RuvC catalytic triad is also mediated by the ∼8 Å shift of a lid helix, which contributes to the change in active site solvent exposure (**Fig. 2f**). This is akin to the lid loop or helix that gates active site exposure reported for other Cas12 endonucleases ^23,24^.

Once activated by target binding, the lid of these endonucleases remains ‘open,’ enabling ssDNA cleavage in *trans*. However, in previously reported Cas12a structures, the RuvC active site is somewhat buried due to the presence of the Nuc domain ^25,26^. In structures of catalytically dead Cas12b and Cas12i, bystander ssDNA bound in *trans* is tightly interwoven to sit within the active site ^22,23^ (**Extended Data Fig. 6**). This is in stark contrast to the highly accessible Cas12a2 RuvC active site in the ternary complex, providing a structural basis for efficient cleavage of a wide range of substrates in *trans*. The lack of a Nuc domain and the presence of a highly exposed RuvC active site in the ternary structure thus explain why Cas12a2 collateral nuclease activation results in an Abi phenotype (Dmytrenko 2022) while Cas12a collateral ssDNase activity does not play a role in bacterial immunity ^27^.

Unlike in Cas12a, the path followed by the target RNA – crRNA duplex completely circumvents the RuvC active site, suggesting that RNA degradation is predominantly in *trans*. To test this, we incubated the Cas12a2 binary complex with fluorescently labeled target RNA at a range of molar ratios and analyzed RNA cleavage. The 5’ FAM label was consistently trimmed due to the ∼20-nt flexible RNA sequence extending from the spacer (**Fig. 2g**). Sybr-gold staining revealed that even though the extended single-stranded 3’ end of the target RNA was trimmed, the target RNA otherwise remained intact and was protected from degradation with a molar equivalence or excess of Cas12a2. This is distinct from the cleavage mechanisms of other Cas12 nucleases that achieve antiphage immunity through cleavage in *cis* ^15,27^, and is reminiscent of Cas13 RNase activity in *cis* and *trans*, where the hybridized region of the target RNA remains intact ^16,17^.

### Collateral dsDNA binding via duplex contortion

We next sought to visualize how Cas12a2 can accommodate and degrade nucleic acid duplexes. To this end, we determined a 2.7 Å-resolution structure of crRNA-guided Cas12a2 bound to both an activating target RNA and a collateral dsDNA substrate analog (**Fig. 3**). The RuvC active site and 11 of 20-bp of the DNA duplex were well resolved, while the flexible DNA ends are only visible at lower density thresholds.

**Fig. 3.**
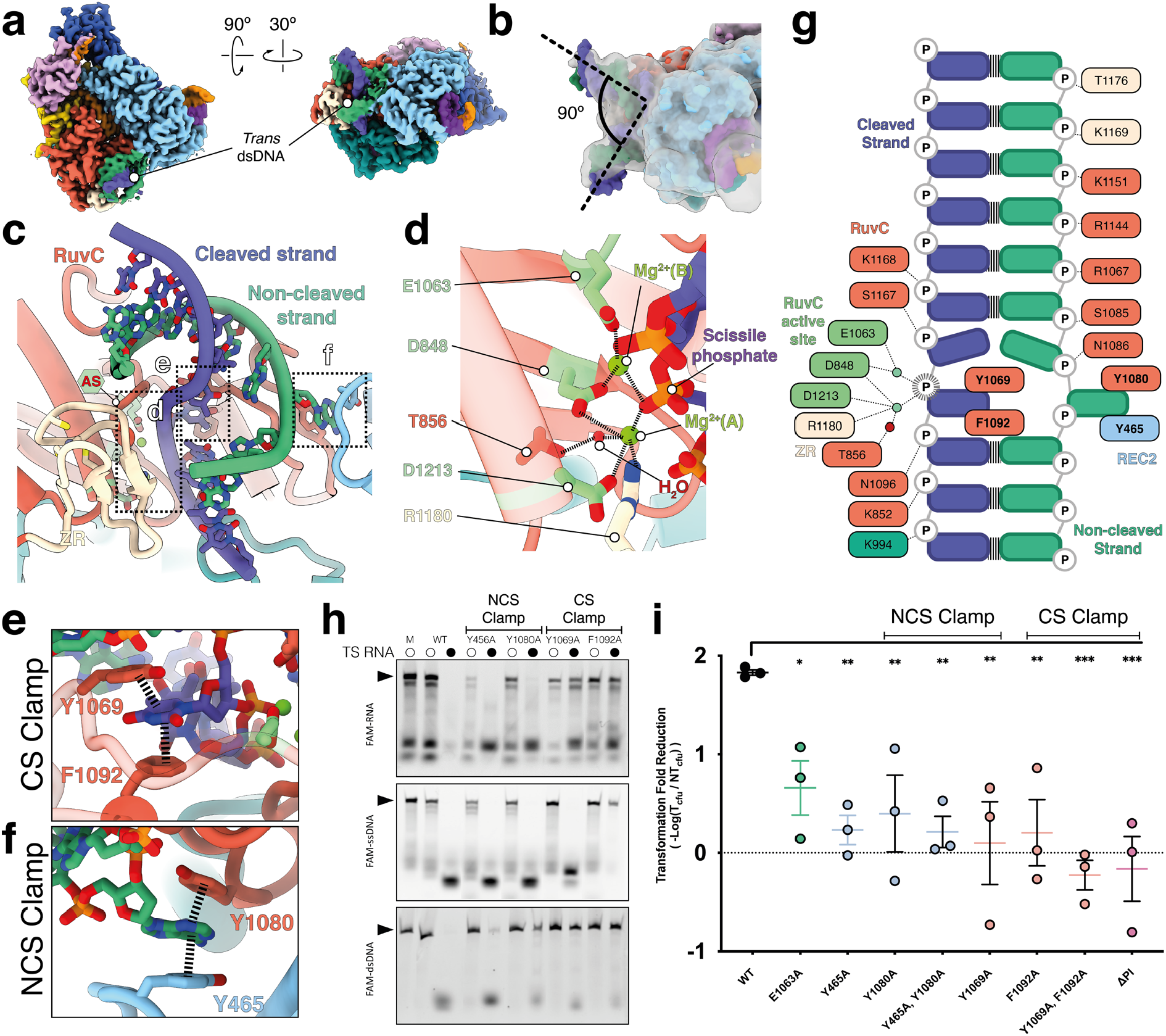
Cas12a2 binds and clamps duplex DNA. **a**, Cryo-EM structure of Cas12a2 quaternary complex. Trans-dsDNA is shown as slate blue and sea green. **b**, Atomic model of Cas12a2 quaternary complex. **c**, dsDNA situated within active site. **d**, Close-up view of Cas12a2 active site. **e**, Cleaved strand held in place through aromatic clamp. **f**, Non-cleaved strand held in place through aromatic clamp. **g**, Schematic of interactions between Cas12a2 and the collateral dsDNA substrate. CS scissile phosphate is denoted by dashed outline. **h**, Cleavage of ssRNA (top), ssDNA (middle) and dsDNA (bottom) by Cas12a2, and aromatic clamp mutants. M – size marker (intact substrate). **i**, Mutations to essential residues resulted in loss of Cas12a2’s ability to clear plasmid (ie. lower transformation fold reduction). Significance between WT and mutant SuCas12a2 was determined by Student T-test. \**P*<0.05, \*\**P*<0.01, \*\*\**P*<0.001.

In our structure, the dsDNA duplex is sharply bent by ∼90º (**Fig. 3b**), resulting in duplex distortion and local melting of two base-pairs in the immediate vicinity of the RuvC active site (**Fig. 3c**), enabling the positioning of the scissile phosphate adjacent to the RuvC catalytic triad (**Fig. 3d**). We designate the DNA strand within the RuvC active site as the cleaved strand (CS), and its complement DNA strand as the non-cleaved strand (NCS) to differentiate from target strand (TS) and non-target strand (NTS) nomenclature used to describe the strands of dsDNA that are specifically targeted with a crRNA guide (e.g. Cas12a or Cas9). As both DNA strands contained non-hydrolyzable phosphothioate modifications, we could visualize the pre-hydrolysis RuvC active site state, including two Mg^2+^ ions and a putative activating water adjacent to one of the ions (designated Mg^2+^(A)) (**Fig. 3d, Extended Data Fig. 7**).

Adjacent to the RuvC active site, both the CS and NCS are stabilized by a large network of non-specific interactions with Cas12a2 (**Fig. 3c,g**). This hub of contacts with both duplex ends induces duplex bending and local melting. The melted bases are subsequently captured by two pairs of ‘aromatic clamps’ (Y465 and Y1080, Y1069 and F1092, respectively) that each hold a single DNA base through π-π stacking, preventing re-hybridization (**Fig. 3e,f**). We confirmed this result through *in vitro* collateral nuclease assays (**Fig. 3h**): while wild-type Cas12a2 degraded ssRNA, ssDNA and dsDNA in *trans* when activated with complementary target RNA, the NCS clamp mutations Y465A and Y1080A reduce and abrogate duplex cleavage, respectively, while having no effect on ssRNase or ssDNase nuclease activity. This indicates that unwinding by the NCS clamp is critical for nuclease activity of a DNA duplex. CS clamp mutations Y1069A and F1092A similarly abrogate dsDNase activity as expected. Intriguingly, while both CS clamp mutations prevent ssRNase activity, only F1092A blocks ssDNase activity (**Fig. 3h**). Since Y1069A preserves ssDNase activity but prevents ssRNase and dsDNase activity, this mutant may enable development of a molecular biosensor that degrades a fluorescence reporter ssDNA upon recognition of a complementary ssRNA, enabling sensitive detection of RNA without target depletion as is the case in Cas13 reporter systems.

As NCS clamp mutations ablate dsDNase activity but have no effect on single-stranded collateral nuclease activity, we tested these separation-of-function mutants *in vivo*. As expected, CS mutations and PI domain truncation blocked Cas12a2 activity (**Fig. 3i**). Mutation of NCS clamp residues individually, and simultaneously, abrogated the ability of Cas12a2 to clear target plasmid *in vivo*, providing direct evidence that aromatic clamp-mediated duplex melting is essential for Cas12a2 activity. These data also suggest that duplex degradation is the driving force behind Cas12a2-mediated immunity, as mutants that retained ssRNase and ssDNase activities were not sufficient to provide immunity.

Collectively these data show that Cas12a2 mediates Abi through a unique mechanism of dsDNA cleavage, distinct from Abi-mediated indiscriminate RNA cleavage seen in other RNA targeting CRISPR-Cas systems ^10,32^. The catalytic mechanism of Cas12a2 is consistent with that of other RuvC endonucleases ^21,30,33,34^, where protein-induced structural tension of the DNA facilitates proper scissile phosphate coordination. However, within the available Cas9 and Cas12a2 structures, the NCS aromatic clamps provide a unique strategy to cleave duplexed nucleic acids.

## Discussion

Our results support a detailed mechanism of Cas12a2 in anti-phage defense. Hybridization of the crRNA to PFS-containing RNA targets drives major conformational changes in Cas12a2, exposing the RuvC active site and alleviating autoinhibition (**Fig. 4**). In the active complex, the RuvC domain is located within a ∼30 Å wide positively-charged groove that is sufficiently large to accommodate duplexed nucleic acids. Non-specific electrostatic interactions facilitate collateral substrate capture, accompanied by duplex distortion and local base-pair melting. The melted bases are stabilized by two pairs of aromatic clamps that enable appropriate positioning of single nucleic acid strands within the RuvC active site. This multiple-turnover DNA nicking culminates in dsDNA degradation, distinct from the single-turnover dsDNA cleavage of Cas12a or Cas9 ^15,35^and enables robust and widespread DNA destruction *in vivo*.

**Fig. 4.**
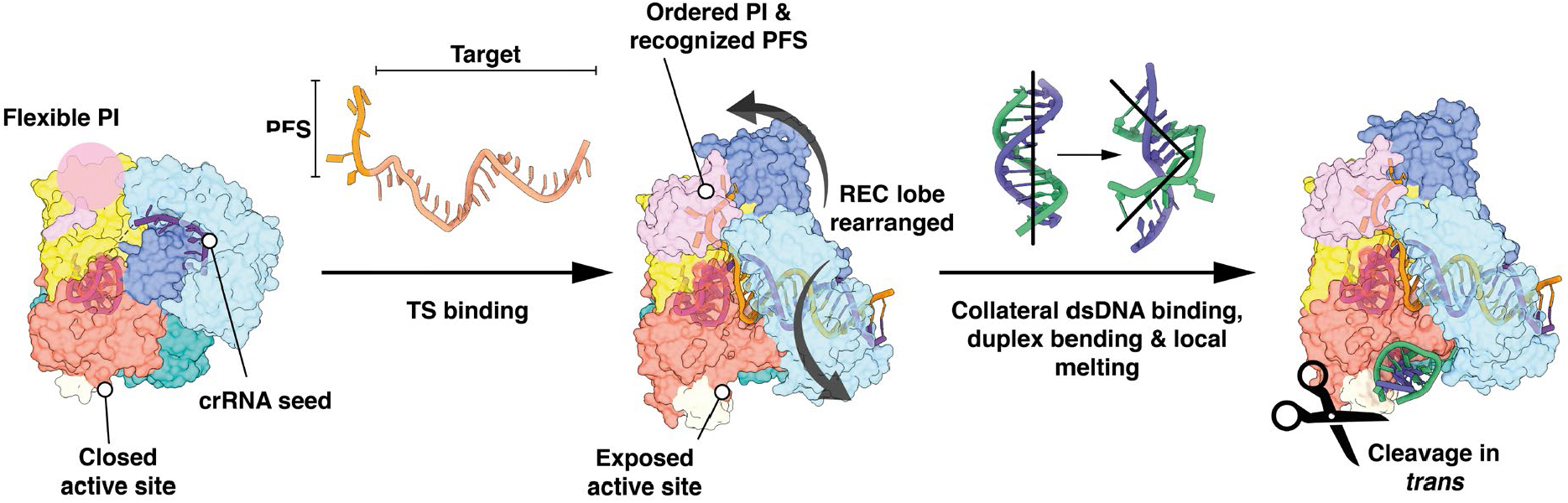
Mechanism of RNA target-activated collateral nuclease activity in Cas12a2. In the binary complex (Cas12a2 – crRNA), the active site is occluded. The 3’ end of the crRNA is a pre-ordered seed, and the PI domain is flexible. Binding of a complementary RNA target strand (TS) containing a protospacer flanking sequence (PFS) triggers significant conformational changes within the REC lobe, exposing the RuvC active site. This enables binding of dsDNA in *trans*, resulting in duplex kinking and local melting. The duplex is held open through the action of two pairs of aromatic clamp residues (**Fig. 3**), enabling nicking by RuvC.

While Cas12a and Cas12a2 can utilize the same crRNA due to their similar WED and RuvC domains, multiple structural divergences confer disparate target preferences, biochemical activities, and enable Cas12a2 to evade inhibition by multiple Cas12a-targetting anti-CRISPR proteins. Notably, the lack of a Nuc domain (also referred to as Target Loading Domain, or Target Nucleic acid Binding domain ^36,37^) may contribute to the inability of Cas12a2 to directly bind dsDNA targets via the formation of an R-loop, conferring specificity for ssRNA target binding. Additionally, the lack of a Cas12a2 Nuc domain increases the accessibility to the RuvC nuclease active site, enabling rapid duplex capture and cleavage in *trans*. Previous structures of Cas12b and Cas12i bound to bystander ssDNA have shown that *trans* ssDNA substrates ^22,23^ must follow a tortuous, narrow path to reach the RuvC active site, which is incompatible with rigid dsDNA substrates (**Extended data Fig. 6**). This is in stark contrast to the highly exposed Cas12 RuvC active site, providing a structural mechanism for dsDNA cleavage in *trans*.

While the mechanism of Cas12a2 is somewhat reminiscent of RNA-targeting by Cas13, there are several distinct differences to Cas12a2. Target binding by Cas13 activates non-specific RNA degradation both in *cis* and in *trans*, resulting in persistent nuclease activation as the phage genome continues to produce target transcripts ^29,32,38^. RNA cleavage can occur in *cis* if the target is of suitable length to fully hybridize with the crRNA and reach the distal active site opposite the seed region ^28^. Similarly, Cas12a2 can trim target RNA in *cis*, but the hybridized target strand remains intact, enabling uninterrupted Cas12a2 activation. Cas13 systems therefore act as sentinels for viral RNA, whereas Cas12a2 represents the nuclear launch button, conferring population-level anti-phage defense through abortive infection.

Our discovery of the Cas12a2 Y1069A point mutant that can cleave ssDNA but not dsDNA or ssRNA provides a blueprint for an RNA-sensor with collateral activity that does not destroy the RNA target it has been programmed to detect, which might enable the development of a highly sensitive RNA diagnostic.

## Methods

### Mutagenesis

The mutant constructs used in this paper were prepared by point directed mutagenesis using the Q5 Site-Directed Mutagenesis Kit (NEB). Primers were designed and annealing temperatures determined using the NEBase Changer tool. All primers were ordered through IDT. Plasmid ligation was achieved with KLD Enzyme Mix (NEB) and completed plasmids were sequenced to verify correct mutations (Plasmidsaurus).

### Expression and Purification

N-terminal hexa-his tagged Cas12a2 and various mutant plasmids were transformed into chemically competent E. coli NiCo21 cells (NEB). A single colony from transformation was selected for starter culture in 20mL LB media grown at 37°C overnight (16 - 18H). Each starter culture was used to inoculate 1.0L of TB media and then cultures were grown up to OD_600_ = 0.6 at 37°C. Cultures were cooled in ice for 15min before being grown at 18°C for an additional 16 - 18H followed by harvesting by centrifugation. Cell pellets were stored at -80°C or used immediately for protein purification.

Cell pellets were resuspended in Lysis Buffer (25mM Tris pH 7.2, 500mM NaCl, 2mM MgCl2, 10mM imidazole, 10% glycerol) treated with protease inhibitors (2 μg/mL Aprotinin, 10 μM Leupeptin, 0.2 mM AEBSF, 1.0 μg/mL Pepstatin) and 1 mg/mL lysozyme and incubated for 30min on ice with shaking. Cells were lysed by sonication and clarified by centrifugation. Clarified lysate was batchbound to nickel resin for 30min at 4°C then allowed to flow through. Lysate was then passed again over nickel resin twice. Nickel resin was washed with Nickel Wash Buffer (25mM Tris pH 7.2, 2M NaCl, 2mM MgCl_2_, 10mM imidazole, 10% glycerol) and eluted with Nickel Elution Buffer (25mM Tris pH 7.2, 500mM NaCl, 2mM MgCl_2_, 250mM imidazole, 10% glycerol). Nickel elutions were desalted into Low Salt Buffer (25mM Tris pH 7.2, 50mM NaCl, 2mM MgCl_2_, 10% glycerol) using either HiPrep 26/10 Desalting column (Cytiva) or Pd10 Sephadex G25M Columns (Cytiva) depending on purification size. Protein samples were then loaded over ion exchange (HiTrap SP HP or HiTrap Q HP column (Cytiva) depending on the isoelectric point of the construct) using Low Salt Buffer and eluted with a gradient of High Salt Buffer (25mM Tris pH 7.2, 1.0M NaCl, 2mM MgCl_2_, 10% glycerol).

Peak elutions were then pooled and concentrated to ∼1.0mL. During this process, protein to be used for biochemistry was desalted by refilling the concentrator with Low Salt Buffer twice to exchange out the High Salt Buffer. Concentrated protein for Cryo-EM was loaded over a HiLoad 26/600 Superdex 200 pg column (GE Healthcare) using SEC Buffer (25mM HEPES pH 7.2, 150mM NaCl, 2mM MgCl_2_, 5% glycerol). Peak fractions were then pooled and concentrated again. Concentrated protein was either flash frozen in liquid nitrogen or used in complex formation for Cryo-EM.

### Complex Formation for Cryo-EM

Before crRNA was added to Cas12a2, RNA was incubated at 65°C for 3 min followed by cooling 1 °C/min to room temperature. Binary complex was formed for Cryo-EM by combining protein and synthetic crRNA in a 1:1.2 molar ratio in SEC buffer (25mM HEPES pH 7.2, 150mM NaCl, 2mM MgCl_2_, 5% glycerol) and incubating at 24°C for 10 min. Unbound crRNA was then separated from the binary complex over a Superdex 200 10/300 increase GL sizing column(Cytiva) into Cryo-EM Buffer (12.5mM HEPES pH 7.2, 150mM NaCl, 2mM MgCl_2_) Eluted protein was concentrated to 30μM in a 100kDa MWCO spin concentrator (Corning) and flash frozen in liquid nitrogen.

### Far-UV Circular Dichroism Spectroscopy

Protein samples of Cas12a2 mutants were prepared at a concentration between 0.3 - 0.5 mg/mL determined by nanodrop in CD Buffer (20 mM Tris pH 7.2, 100 mM NaCl). Far-UV Circular dichroism (CD) readings utilized a Jasco-J1500 spectropolarimeter (Easton, MD). The CD spectra were obtained from 260–190 nm using a scanning speed of 50 nm/min (with a 2 s response time and accumulation of three scans). Melting curves of ΔPI Cas12a2 samples (in sealed quartz cuvettes with 0.1 cm pathlength) were obtained by monitoring the CD signal at 222 nm every 1°C over a 10–90°C temperature range, employing a temperature ramp of 15°C/h. The CD signal was converted to molar ellipticity by Jasco Spectra Manager software.

### Plasmid Curing Assay

Plasmid curing assays were conducted as described previously (Dmytrenko et al., 2022). In short, Immune system plasmids were prepared with Cas12a2 and a 3x CRISPR repeat and transformed into BL21 AI cells by heat shock transformation. Cells expressing the immune system were then made electrocompetent (J. Sambrook, D.W. Russell, Molecular Cloning: A Laboratory Manual (2001).) and immediately transformed by electroporation with 50ng of either target or non-target plasmid. Transformations were recovered for 18 H in 450 uL LB media containing 1 mM IPTG, 0.2% L-arabinose and antibiotics for the immune system plasmid. Recovered transformations were then serially diluted in LB media between 10^1^ and 10^6^ and spotted in 10 μL drops on LB Agar plates containing 1 mM IPTG, 0.2% L-arabinose and antibiotics for both immune system and target or non-target plasmids. Colonies were counted in the highest countable spot in the dilution series and the relative transformation efficiency was calculated between the target and non-target plasmid.

### Activation Assay

Binary complex of either WT or ΔPI Cas12a2 with crRNA was combined with various targets (FL, -PFS, -5, -10) to final reaction conditions of 600 nM Cas12a2, 720 nM crRNA and 300nM FAM labeled target in 1x NEB 3.1 Buffer (50mM Tris pH 7.9, 100mM NaCl, 10mM MgCl_2_, 100μg/mL BSA). Binary complex was first formed by combining WT or ΔPI Cas12a2 with crRNA in a 1:1.2 molar ratio and incubating 30 min at room temperature with NEB 3.1 buffer as a 2x master mix. Binary complex and target RNAs were then combined to their final reaction concentration and incubated at room temperature for 1.0 H before quenching with 1:1 (v/v) phenol-chloroform pH 4.5 mixed by flicking.

Reactions were quenched with phenol-chloroform pH 4.5 and mixed by flicking followed by spinning down for 30 sec. Reaction products were run on a 12% FDF page gel as described by Harris and co. (C. J. Harris, A. Molnar, S. Y. Müller, and D. C. Baulcombe, “FDF-PAGE: A powerful technique revealing previously undetected small RNAs sequestered by complementary transcripts,” Nucleic Acids Res., vol. 43, no. 15, pp. 7590–7599, 2015.) with minor modifications. Loading dye was replaced with 30% glycerol and gels were run at 50 V for 15 min before increasing voltage to 150 V.

### Target Protection Assay

Binary complex of Cas12a2 and crRNA was combined with FAM labeled FL target RNA to a final reaction condition of 600 nM Cas12a2, 720 nM crRNA, 300nM Target RNA in 1x NEB 3.1 Buffer (50mM Tris pH 7.9, 100mM NaCl, 10mM MgCl_2_, 100μg/mL BSA). Binary complex was first formed as described in the activation assay. The 2x master mix was then combined with the target RNA and incubated at 37°C, with time points (5, 15, 30, 60 and 120 min) extracted and quenched with phenol-chloroform pH 4.5. Samples were then run on an 12% FDF page gel as described in the activation assay. Completed gels were imaged for FAM fluorescence and then stained with Sybr Gold and imaged again to show unlabeled RNA species.

### Concentration Dependent Effect on Cleavage

FL FAM labeled Target RNA (300nM) was combined with a range of binary complex concentrations (600, 300, 150, 75, 37.5 nM) to achieve Complex: Target ratios of 2:1, 1:1, 1:2, 1:4, and 1:8. Binary complex was first formed by combining Cas12a2 (1200 nM) with crRNA in a 1:1.2 molar ratio and incubating 30 min at 37°C with NEB 3.1 buffer as a 2x master mix. Binary complex was then serially diluted in 2x NEB 3.1 Buffer (100mM Tris pH 7.9, 200mM NaCl, 20mM MgCl_2_, 200μg/mL BSA) to form a 2x master mixes for each complex:target ratio. Each 2x master mix was combined with target RNA and incubated at 37°C for 1.0 H followed by quenching with phenol-chloroform pH 4.5. Quenched samples were visualized as described in the activation assay.

### Trans-Cleavage Assay

Reactions containing 600 nM Cas12a2 (WT or mutants) and 720 nM crRNA were combined with 300 nM FL target RNA and 300 nM FAM labeled non-target RNA, ssDNA or dsDNA. Binary complex was first formed as described previously as a 4x master mix. The master mix was combined with target and non-target substrates and incubated at 37°C for 1.0 H and quenched with phenol-chloroform pH 4.5. Samples were then separated on an 12% 7M Urea page gel and imaged for FAM fluorescence.

### Cryo-EM sample preparation, data acquisition, and processing

Flash-frozen Cas12a2 binary complex was rapidly thawed. 4 µL of the binary complex (XµM) was applied to C-flat holey carbon grids (2/2, 400 mesh) which had been plasma cleaned for 30 seconds in a Solarus 950 plasma cleaner (Gatan) with a 4:1 ratio of O_2_/H_2_. Grids were blotted with Vitrobot Mark IV (Thermo Fisher) for 2 seconds, blot force 4 at 4ºC & 100% humidty, and plunge-frozen in liquid ethane. Data were collected on a FEI Glacios cryo-TEM equipped with a Falcon 4 detector. Data was collected in SerialEM, with a pixel size of 0.94 Å, a defocus range of -1.5 - -2.5 µm, and a total exposure time of 15s resulting in a total accumulated dose of 40 e/Å^2^ which was split into 60 EER fractions. Motion correction, CTF estimation and particle picking was performed on-the-fly using cryoSPARC Live v4.0.0-privatebeta.2^39^. 1,577 movies were collected, of which 1,159 were accepted based on meeting the criteria of a CTF fit of 5 Å or better. All subsequent data processing was performed in cryoSPARC v3.2^40^.

From 987,122 particles picked, 214,647 were selected from a single round of 2D classification. These particles were subjected to ab initio reconstruction (3 classes) followed by heterogeneous refinement resulted in a final subset of 97,470 particles that yielded a 3.46Å-resolution structure from non-uniform refinement. Re-extraction of this subset of particles in a 320 pix box size, splitting of particles into 4 exposure groups and performing per-group CTF refinement and per-particle defocus optimization as implemented in non-uniform refinement^41^ resulted in a 3.2 Å-resolution reconstruction that was used for modelling.

For the ternary complex, a rapidly thawed Cas12a2 binary complex fraction was supplemented with 4-fold excess of heat-annealed (90ºC for 5 mins, and rapidly cooled to 4ºC) PFS-containing RNA target and incubated at room temperature (∼25ºC) for 30 mins prior to vitrification, which was performed in an identical manner to the binary complex as described above. Data were collected using a FEI Titan Krios cryo-electron microscope equipped with a K3 Summit direct electron detector (Gatan, Pleasanton, CA). Images were recorded with SerialEM^42^ with a pixel size of 0.81 Å. A total accumulated dose of 70 electrons/Å^2^ during a 6 second exposure was fractionated into 80 frames. A total of 6,940 micrographs were collected, of which 6,614 with CTF fits of 5 Å or better were retained. On-the-fly processing was performed as described above.

A total of 3,515,037 particles were picked, of which 2,212,319 were selected after 2D classification. Multiple rounds of ab initio reconstruction and heterogeneous refinement resulted in a subset of 192,639 particles were reconstructed to 2.92 Å-resolution using non-uniform refinement as described above. This map was then used for modelling

For the quaternary complex, ternary complex was prepared as described above, and incubated with heat-annealed phosphothioate dsDNA duplex for 30 mins at room temperature. 2.5 µl of complex was applied to C-flat grids (1.2/1.3, 300 mesh) and blotted for 6 seconds, blot force 0 at 4ºC and 100% humidity prior to vitrification. Data were collected on a FEI Glacios cryo-TEM equipped with a Falcon 4 detector, as described for the binary complex. 1,755 movies were collected, of which 1,539 had CTF fits of 5 Å or better and were retained for subsequent processing. On-the-fly motion correction, CTF estimation and particle picking were performed as described above.

1,692,368 particles were picked, of which 425,770 were retained after a single found of 2D classification. A single round of ab initio reconstruction (3 classes) followed by heterogeneous refinement yielded a subset of 260,958 particles which were reconstructed to 2.97 Å resolution using non-uniform refinement. Additional rounds of ab initio reconstruction and heterogeneous refinement were used to further classify particles, resulting in a final subset of 104,857 particles. Extraction of said particles with a 384-pixel box, splitting particles into 9 exposure groups, and reconstruction using non-uniform refinement with per-group CTF refinement and per-particle defocus optimization resulted in a 2.74 Å-resolution reconstruction which was then used for modelling.

### Model building and figure preparation

A Cas12a binary complex (PDB 5NG6)^20^ was rigid body fitted into the Cas12a2 binary complex map. While most of the model did not correspond to the map, the RuvC and WED domains generally were consistent. However, pairwise blast of the two proteins revealed significant gaps or inserts for the relative insertions. However, a single 20 residue hairpin of the Cas12a RuvC domain fitted the Cas12a2 map well and had no gaps or insertions in the pairwise blast. This fragment was isolated was rigid body fitted into the Cas12a2 map, and the sequence was mutated to the corresponding region of Cas12a2 using Coot^43^. This was then used as a fiducial to build the rest of the complex de novo using in Coot. Attempts at using AlphaFold2 (AF2)^44^ to generate fragments to fit in the map were unsuccessful since it was not possible since adjacent residues within the WED and RuvC domains were separated by protein sequence and the REC1 and REC2 domain boundaries were not obvious from the sequence alone. However, AF2 was used to validate modelling of small structural domains after-the-fact, where smaller, compact regions of the model were folded using AF2, and fitted into the map, indicating correct modelling. This was particularly useful when the *de novo* model contained gaps due to local flexibility.

The 5’ crRNA handle was built using a the Cas12a 5’ crRNA as a template (PDB 5NG6). In the binary complex, the 7-nt 3’ seed region was modelled *de novo* as polyU since it was not possible to unambiguously identify nucleotide identity.

Once fully modelled, Isolde^45^ was used to improve the fit of the model to the map, and real-space refinement as implemented within Phenix^46^ was performed to optimize model geometry.

For the Cas12a2 ternary complex, the RuvC, WED, and part of the Insert domains were in the same conformation as in the binary complex. These were rigid body fitted into the ternary complex map. The REC1, REC2 and the C-terminal half of the Insertion domain were separately fitted as rigid bodies into the ternary complex map. The PI domain structure was predicted using AF2, and then manually connected to the rest of the model. The crRNA – target RNA duplex was modelled as ideal A-form RNA within Coot, and manually connected to the 5’ crRNA handle. The target RNA 3’PFS was modelled *de novo*. Coot was used to fit in gaps within the model, and Isolde was then used to improve the quality of model prior to real-space refinement as described above.

For the quaternary complex, the ternary complex structure was rigid body fitted into the map, and then flexibly fitted and using Isolde. The ZR domain structure was predicted using AF2, and manually connected to the rest of the model. The dsDNA duplex was modelled de novo, with one strand modelled as polyT and the other as polyA since it was not possible to unambiguously determine nucleotide identity. Mg^2+^ and Zn^2+^ ions and an activating H_2_O were modelled manually using the sharpened map. Isolde and real-space refinement were performed as described above.

All structural figures and movies were generated using ChimeraX^47,48^, apart from the modevectors, which were generated in PyMol. Schematic figure was created with BioRender.com.

## Competing Interests

J.P.K.B, T.H., R.N.J, and D.W.T are inventors on a filed patent application based on this work.

## Author Contributions

J.P.K.B. performed cryo-EM structure determination, model building, and structural analysis. T.H. and B.N. purified and reconstituted the enzyme complexes, T.H. performed biochemical and in vivo experiments. C.L.B. aided with in vivo experiments. J.P.K.B., T.H., R.N.J., and D.W.T. analyzed and interpreted the data and wrote the manuscript. R.N.J. and D.W.T. supervised the study and obtained funding for the work.

## Data Availability

The atomic models of Cas12a2 binary, tertiary and quaternary complexes have been deposited into the Protein Data Bank as PDB 8D49, 8D48 and 8D4A, and the corresponding maps have been deposited to the Electron Microscopy DataBank as EMD-29178, -27180 and -27179, respectively. Requests for materials can be sent to R.N.J. and D.W.T.

## Acknowledgements

We thank members of the Jackson and Taylor lab for helpful discussions. A. Brilot for expert cryo-EM technical assistance. O. Dmytrenko for initial guidance on the in vivo plasmid interference assays. Data were collected at the Sauer Structural Biology Laboratory at the University of Texas at Austin. This work was supported in part by an ERC Consolidator grant (865973 to C.L.B.), National Institute of General Medical Sciences (NIGMS) of the National Institutes of Health (NIH) R35GM138080 (to R.N.J.) and R35GM138348 (to D.W.T.), and a Robert J. Kleberg, Jr. and Helen C. Kleberg Foundation Research Grant (to D.W.T.). D.W.T is a CPRIT Scholar supported by the Cancer Prevention and Research Institute of Texas (RR160088).

---

**Extended Data Fig. 1.**
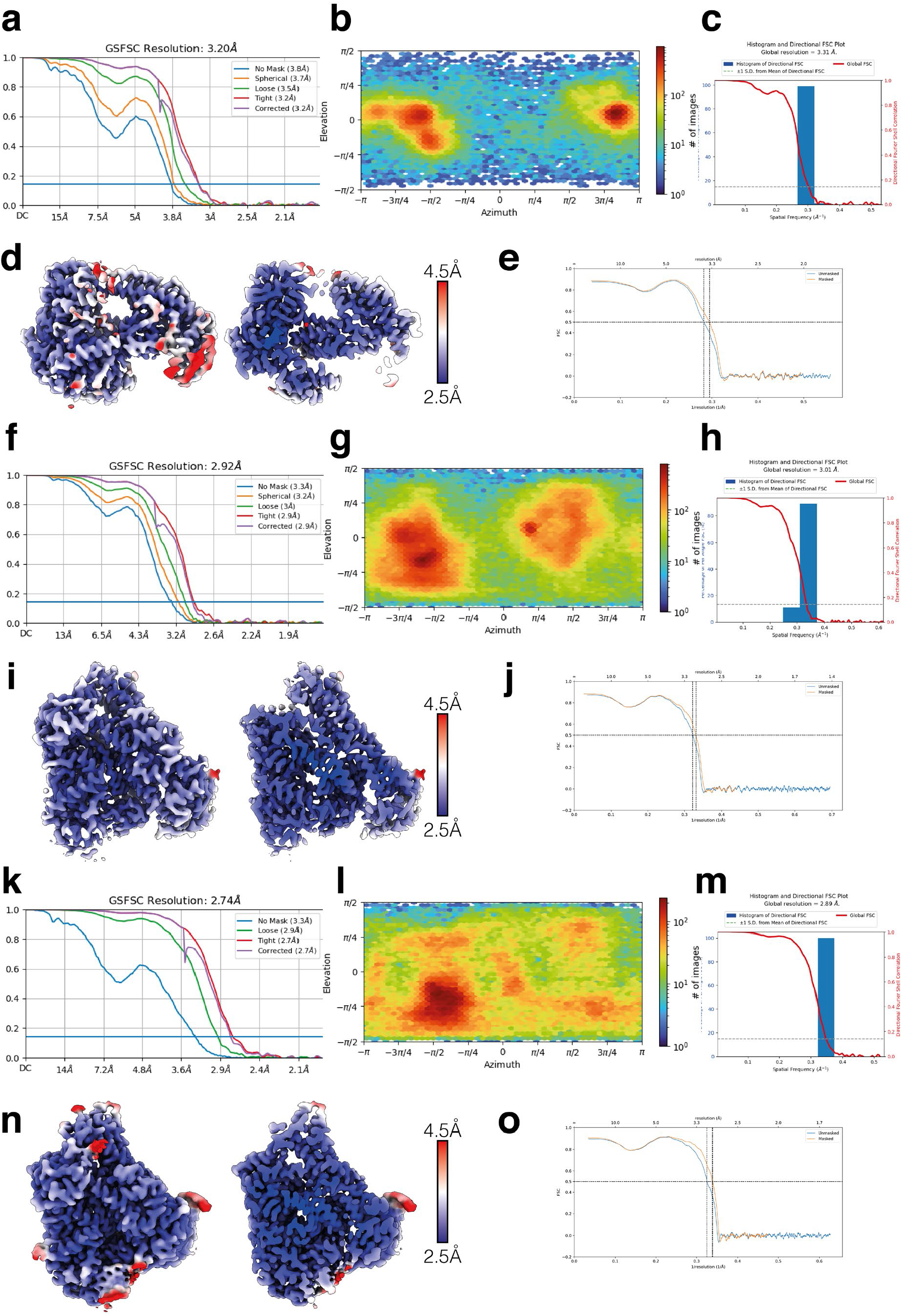
Cryo-EM data analysis. **a**, Gold standard Fourier Shell Correlation (FSC) curves for Cas12a2 binary complex. **b**, Euler plot for Cas12a2 binary complex. **c**, Directional FSC ^49^ for Cas12a2 binary complex. **d**, Cas12a2 binary complex colored by local resolution. Left – surface, right, cut-through. **e**, Map-to-model FSC for Cas12a2 binary complex. **f**, Gold standard FSC for Cas12a2 ternary complex. **g**, Euler plot for Cas12a2 ternary complex. **h**, Directional FSC for Cas12a2 ternary complex. **i**, Cas12a2 ternary complex colored by local resolution. Left – surface, right, cut-through. **j**, Map-to-model FSC for Cas12a2 ternary complex. **k**, Gold standard FSC for Cas12a2 quaternary complex. **l**, Euler plot for Cas12a2 quaternary complex. **m**, Directional FSC for Cas12a2 quaternary complex. **n**, Cas12a2 quaternary complex colored by local resolution. Left – surface, right, cut-through. **o**, Map-to-model FSC for Cas12a2 quaternary complex.

**Extended Data Fig. 2.**
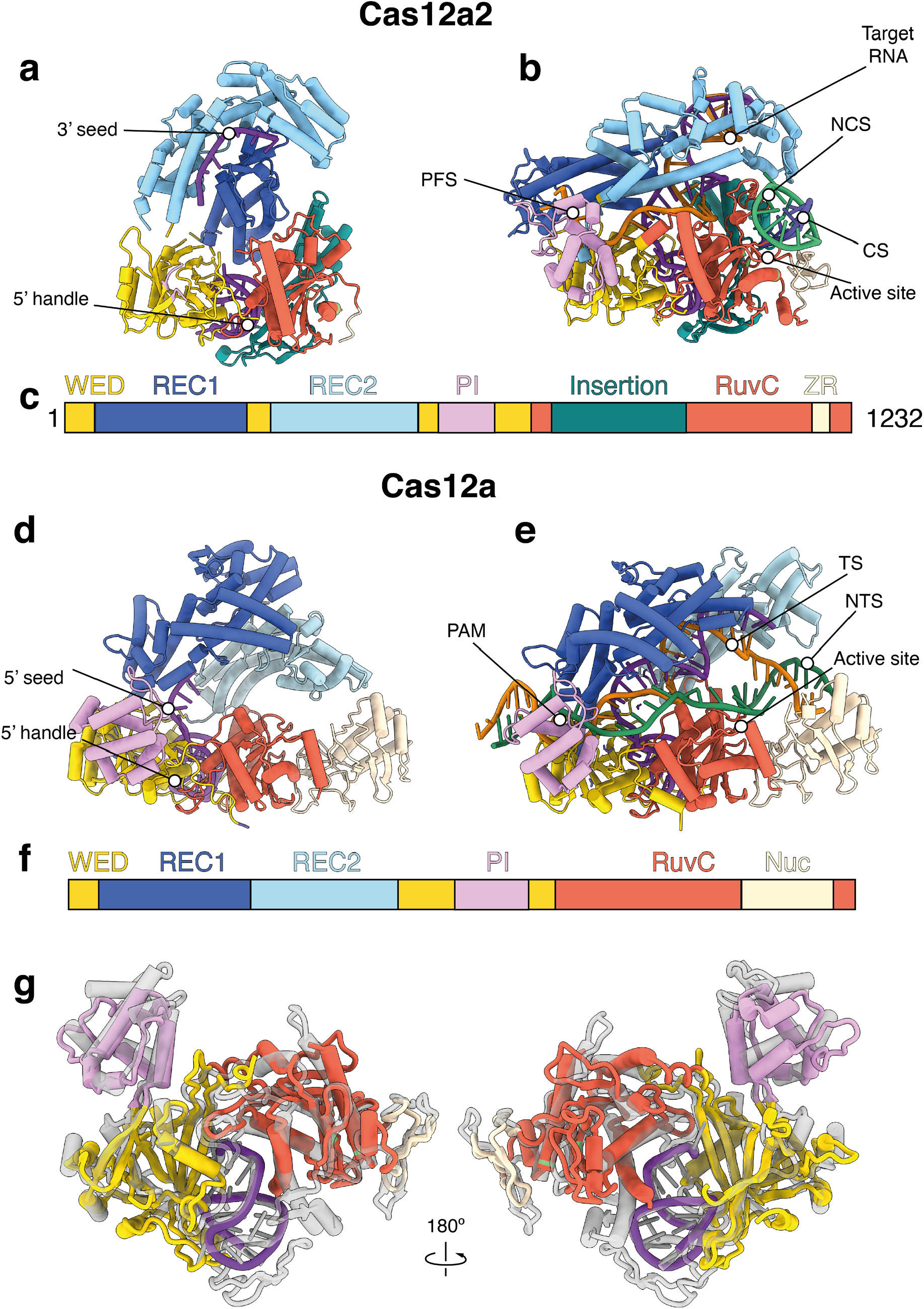
Comparison of Cas12a2 and Cas12a. **a - c**, Cas12a2 binary complex (**a**), active quaternary complex (**b**) colored by structural domain (**c**). **d – f**, Cas12a binary complex (**d**), active ternary complex (**e**) colored by structural domain (**f**). **g**, Overlay of 5’ crRNA handle, WED, RuvC and PI domains of Cas12a2 (colored) and Cas12a (transparent grey).

**Extended Data Fig. 3.**
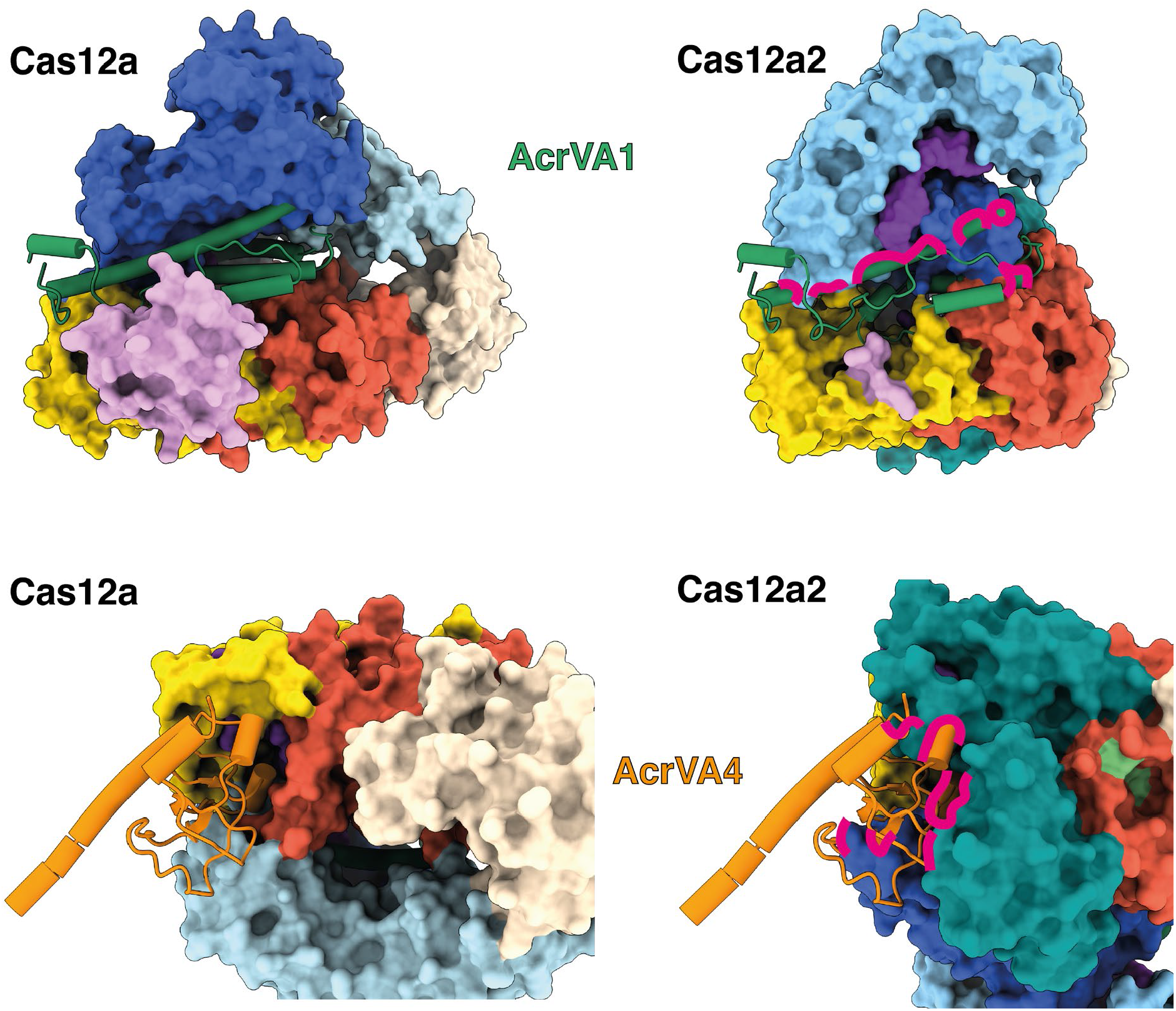
Structural basis for Anti-CRISPR evasion by Cas12a2. Top – Cas12a binary complex (PDB 5NG6, left) and Cas12a2 (right) with AcrVA1 (PDB 6NMD, green). AcrVA1 interacts with the 5’ crRNA seed in Cas12a, which is not present in Cas12a2. There are also significant clashes between AcrVA1 and the REC lobe of Cas12a2. Middle – Cas12a (left) and Cas12a2 (right) binary complex with a monomer of AcrVA4 (PDB 6NMA, orange). There are severe clashes between the Cas12a2 Insertion domain (dark cyan) and AcrVA4. Clashes are shown as magenta lines.

**Extended Data Fig. 4.**
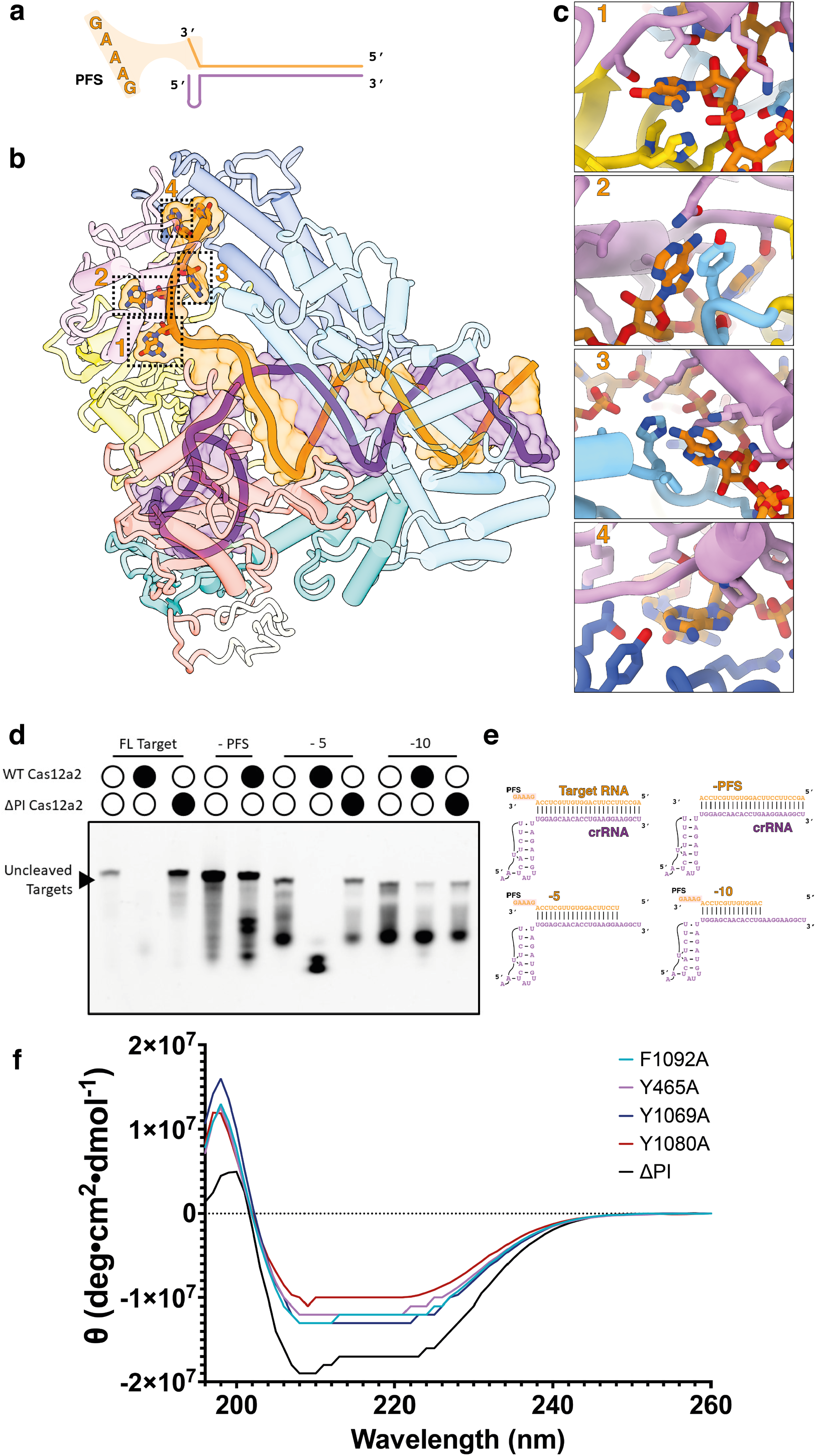
Mechanism of protospacer-flanking sequence (PFS) recognition by Cas12a2. **a**, Schematic representation of PFS (GAAAG) located at the 3’ end of the RNA target RNA. **b**, Positions of the PFS within the Cas12a2 active ternary complex. **c**, Zoom-in of interactions between Cas12a2 and the first four bases of PFS. **d**, Activation of Cas12a2 or Cas12a2 ΔPI cleavage by truncated target RNAs. **e**, Circular Dichroism (CD) spectroscopy of Cas12a2 mutants, including ΔPI truncation. All mutants are properly folded.

**Extended Data Fig. 5.**
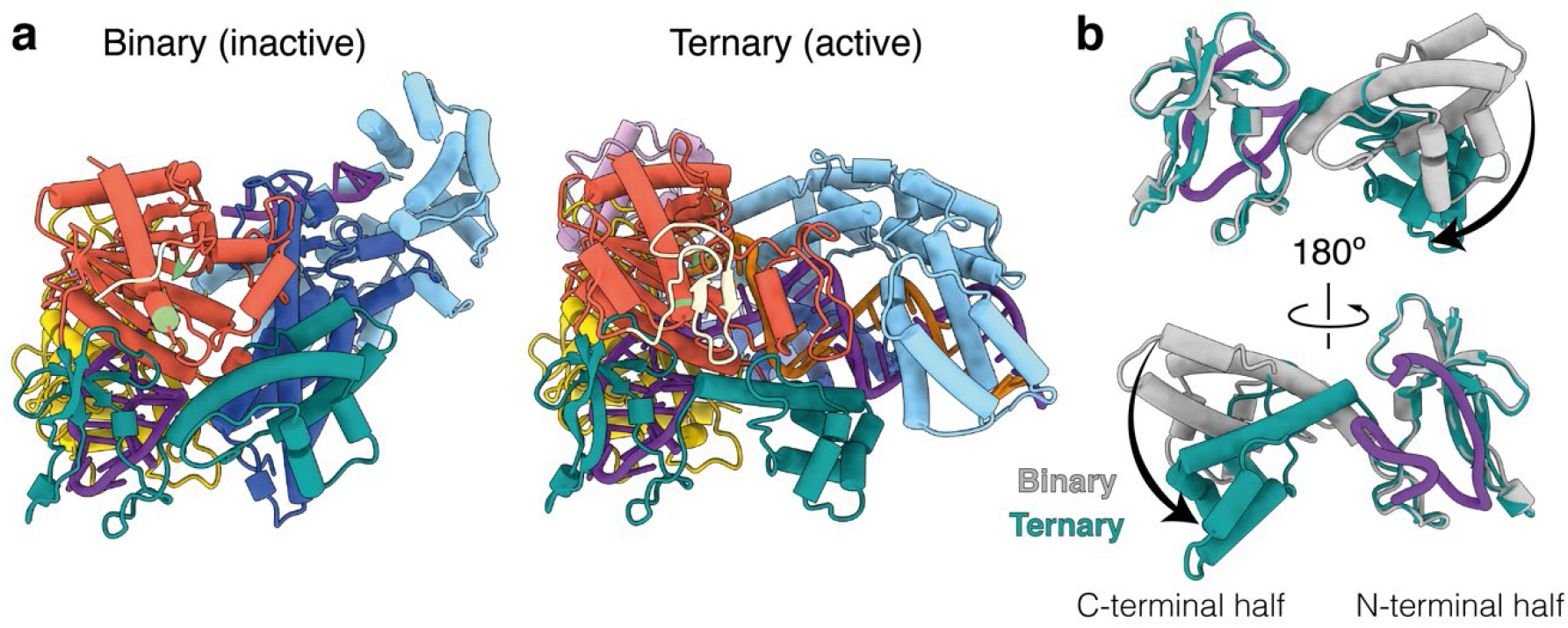
Conformational changes in Insertion domain upon target RNA binding. **a**, Cas12a2 autoinhibited binary complex (left) and active ternary complex (right), colored as in **Fig. 1a**. Insertion domain is dark cyan. **b**, conformational changes of the C-terminal half of insertion domain from binary (grey) to ternary (dark cyan) complexes. N-terminal half remains static, and interacts with the crRNA 5’ handle.

**Extended Data Fig. 6.**
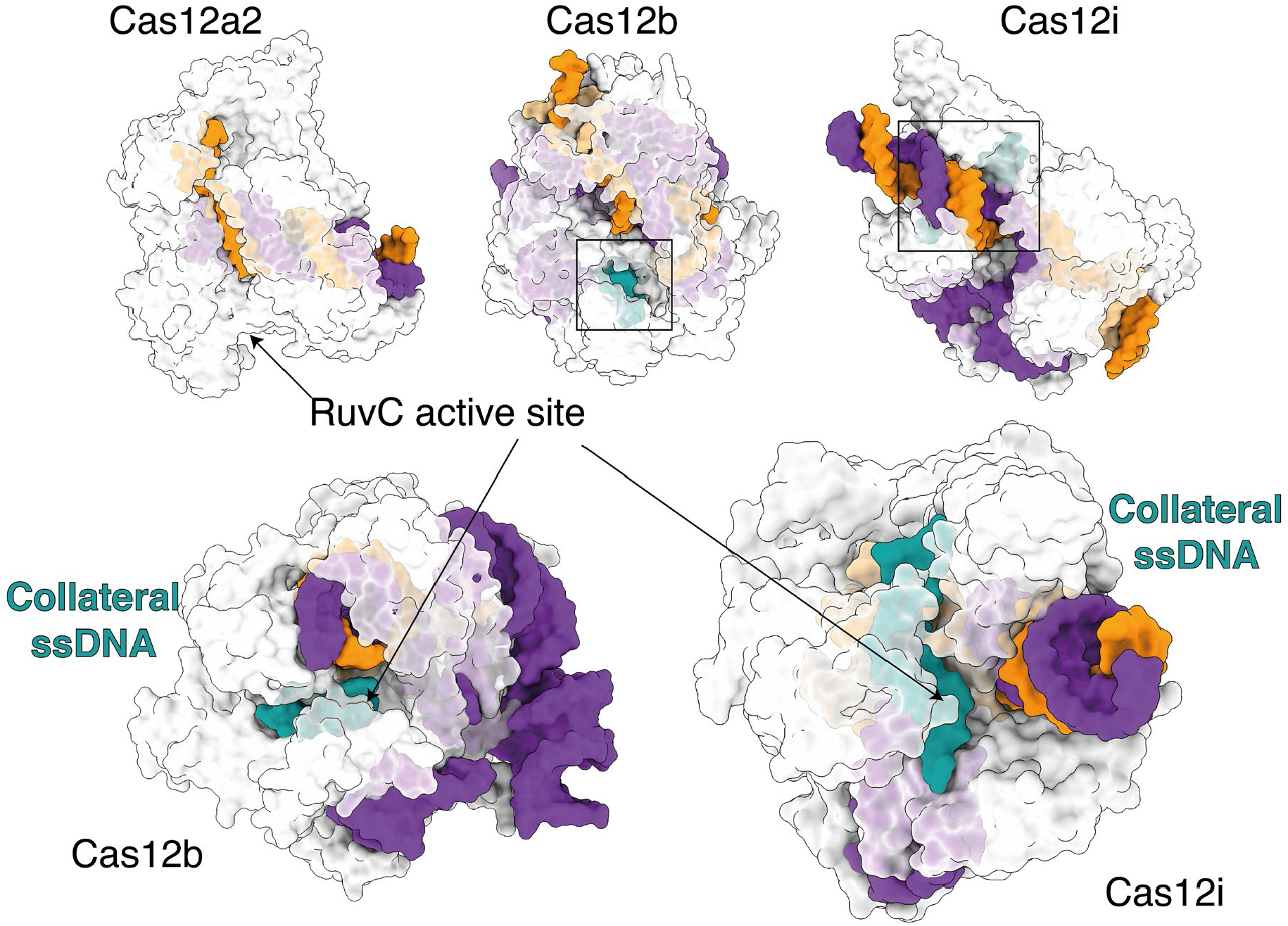
Collateral ssDNA in other Cas12 structures. **Top**, Cas12a2 ternary complex (this study), Cas12b (PDB 5U31) and Cas12i (6W5C). Arrow denotes Cas12a2 active site. Boxes show position of bound collateral ssDNA in Cas12b and Cas12i. **Bottom**, Zoom-in of collateral ssDNA (dark cyan) buried in RuvC active site of Cas12b and Cas12i.

**Extended Data Fig. 7.**
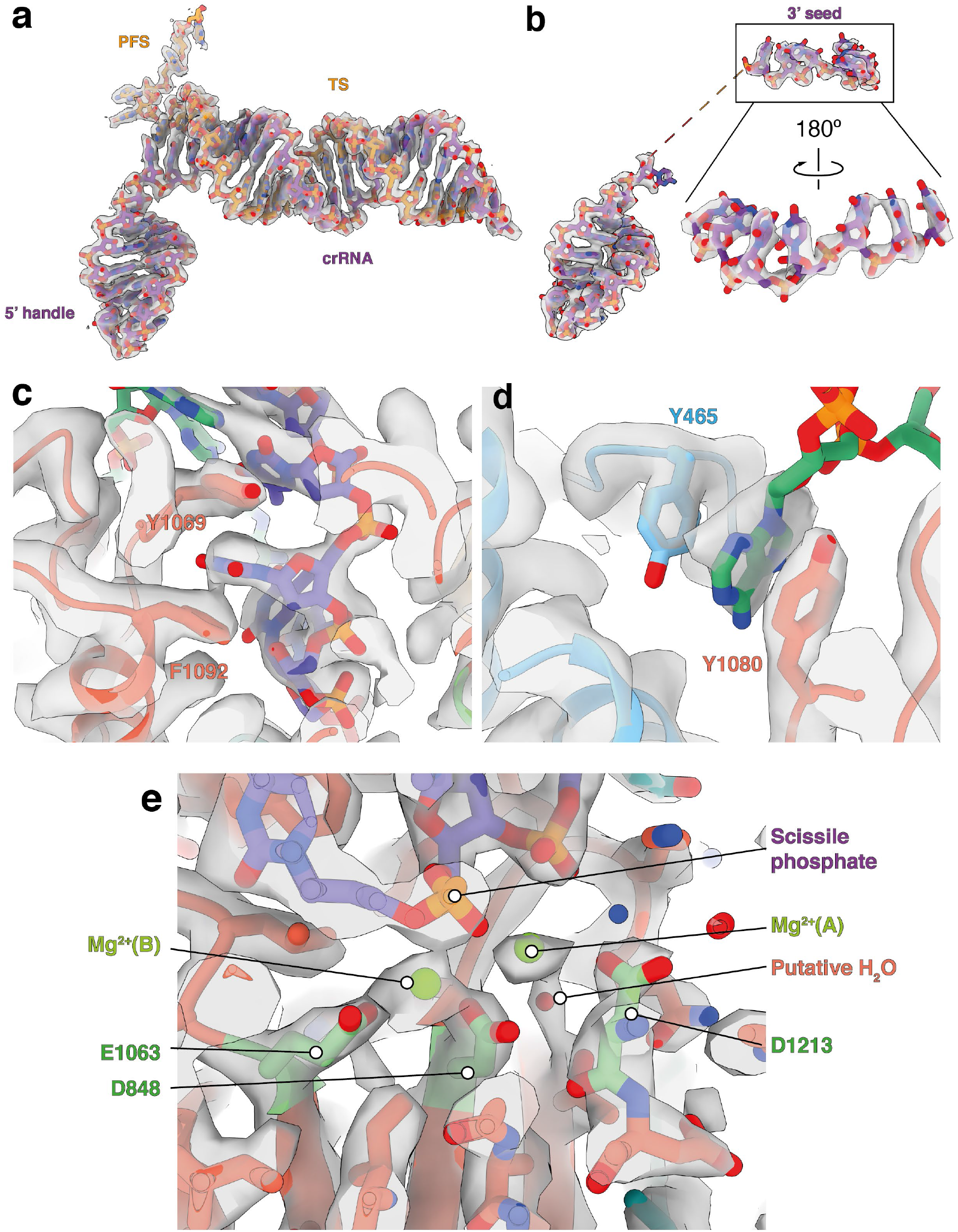
Representative cryo-EM densities. **a, target RNA** and crRNA from ternary complex map, showing 5’ crRNA handle and 3’ target RNA PFS. **b**, 5’ handle and 7-nt pre-ordered seed from Cas12a2 binary complex. **c**, Cleaved strand pair of aromatic clamps. **d**, non-cleaved strand (NCS) aromatic clamps. Density for the unwound bases of the NCS is diffuse due to flexibility, but the nucleobase held within the aromatic clamp is well-resolved. **e**, RuvC active site, including density for scissile phosphate, two Mg^2+^ ions and a putative activating water.

**Extended Data Fig. 8.**
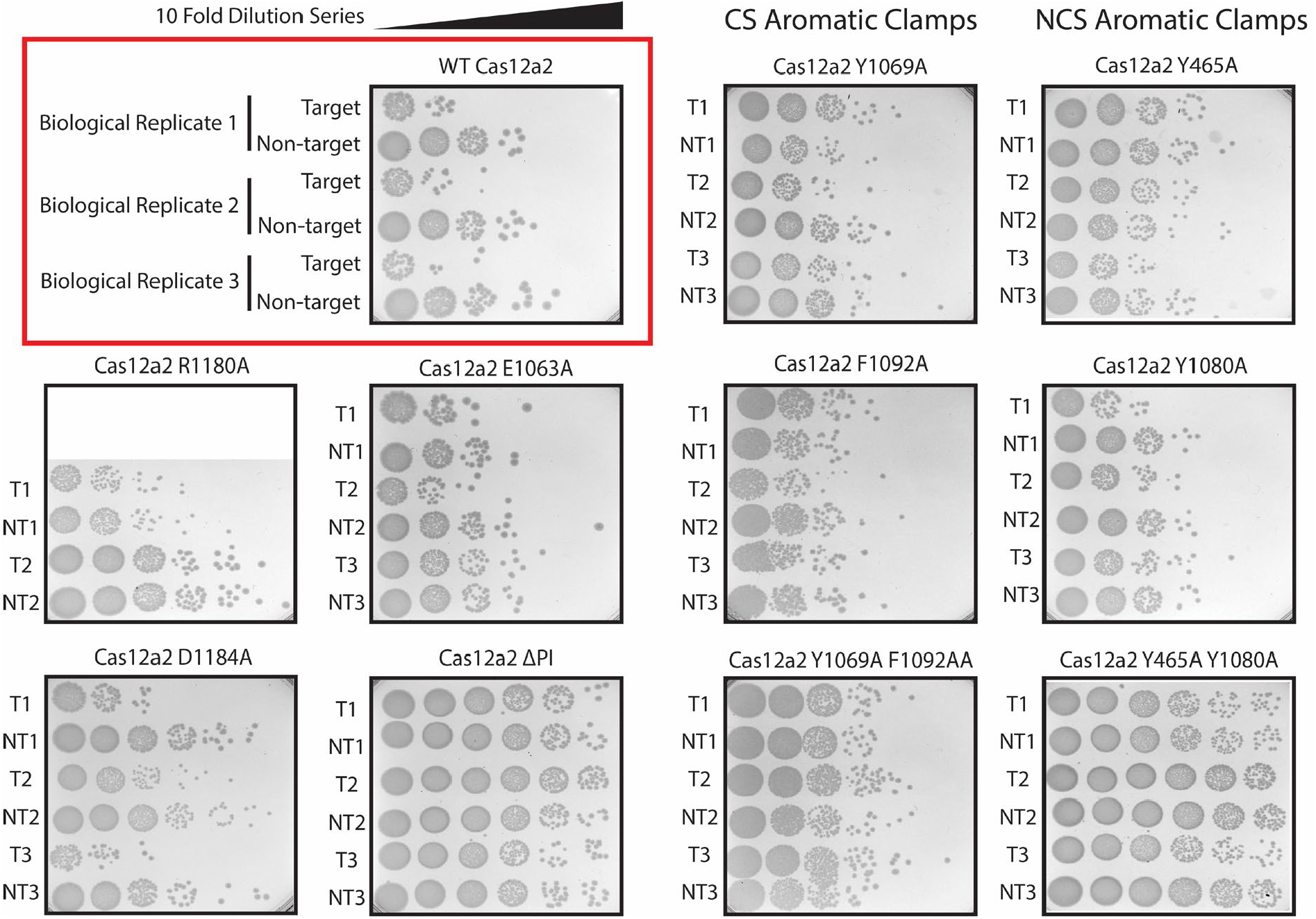
In vivo interference assays. Transformation efficiency of Target (T) and Non-Target (NT) plasmids into electrocompetent *E. coli* containing WT or mutant Cas12a2+3xcrRNA immune system plasmids shown by spot assay. Transformations were plated as 10x serial dilutions between 10^1^ and 10^6^ and the number of colonies in the highest countable spot were used to determine the total number colony forming units (cfu) per 50 ng plasmid. Transformation fold reduction was calculated as -Log_10_(T_cfu_/NT_cfu_). Active immune systems present a ≥100-fold difference between T and NT transformation efficiency. All transformations were completed to at least biological duplicate.

**Extended Data Table 1.**
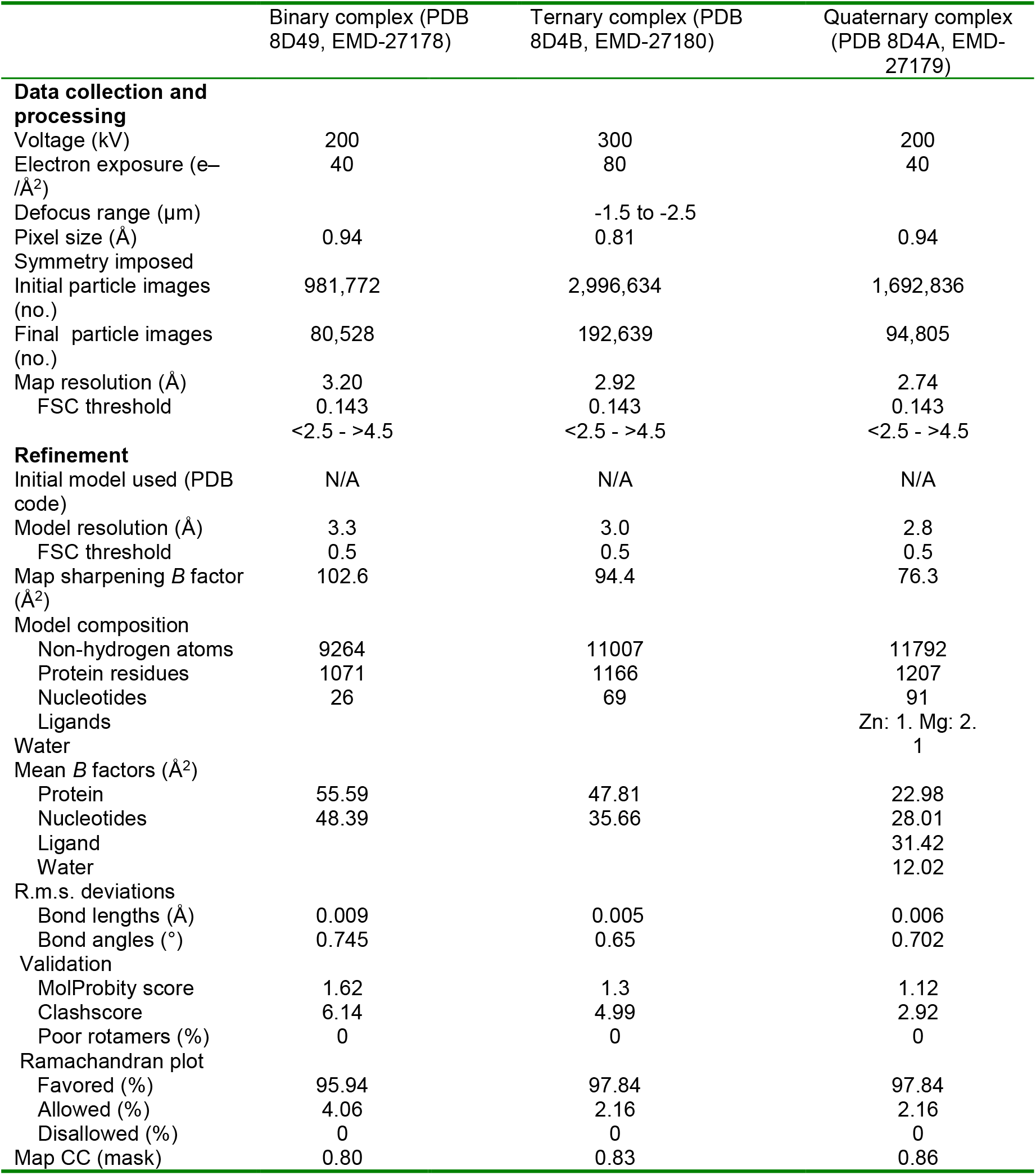
Cryo-EM data collection and model validation statistics.

**Extended Data Table 2.**
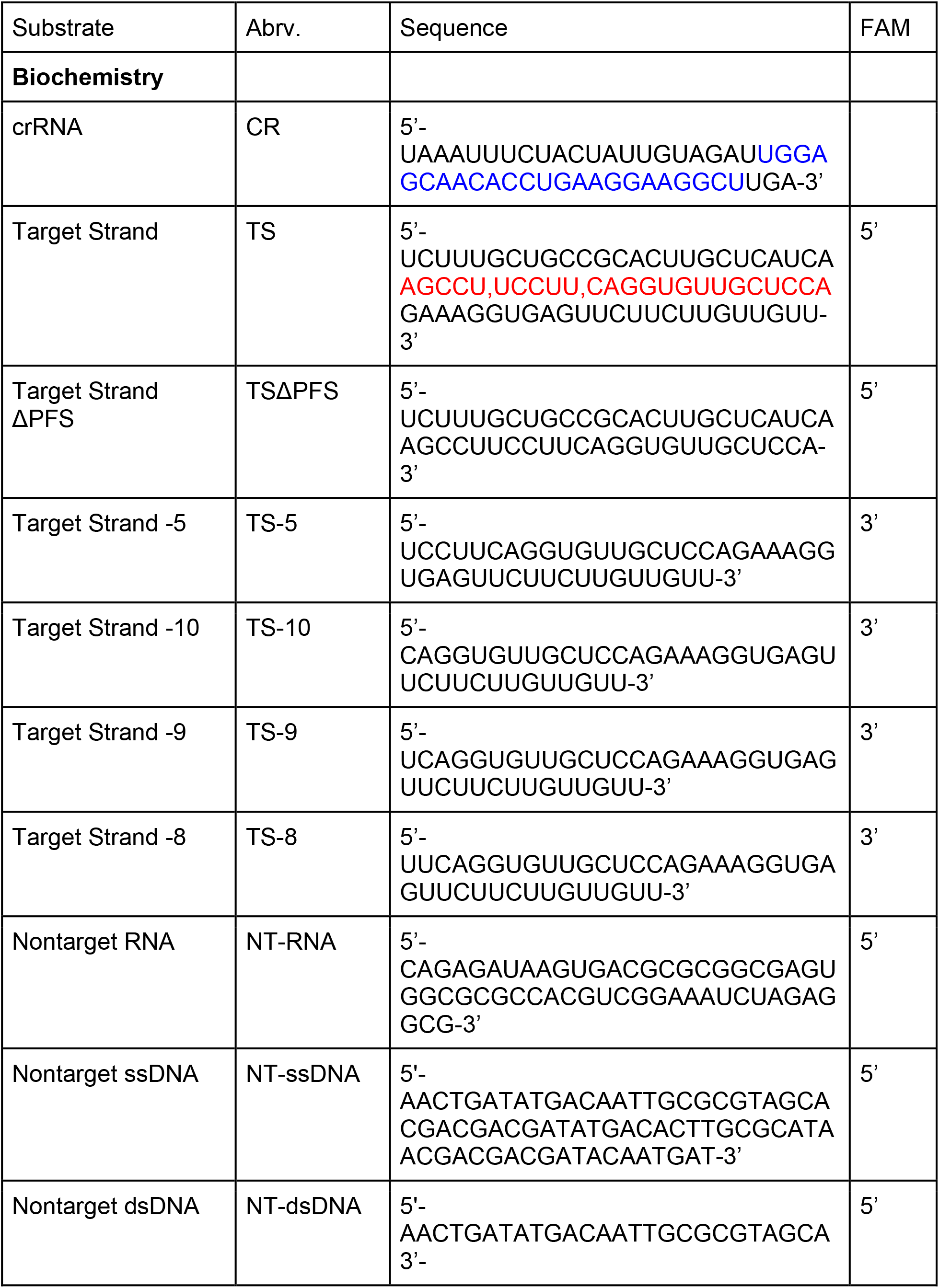
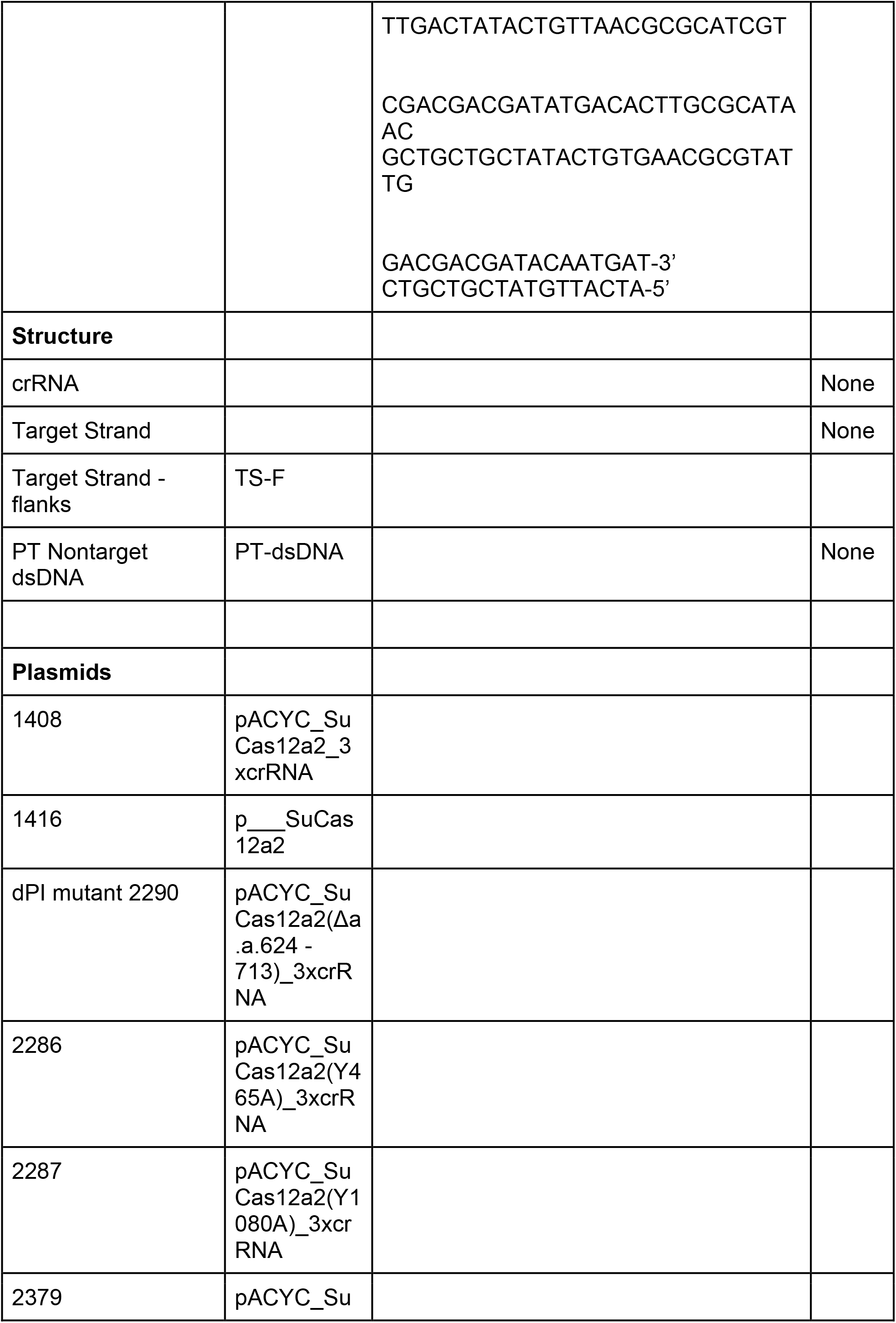
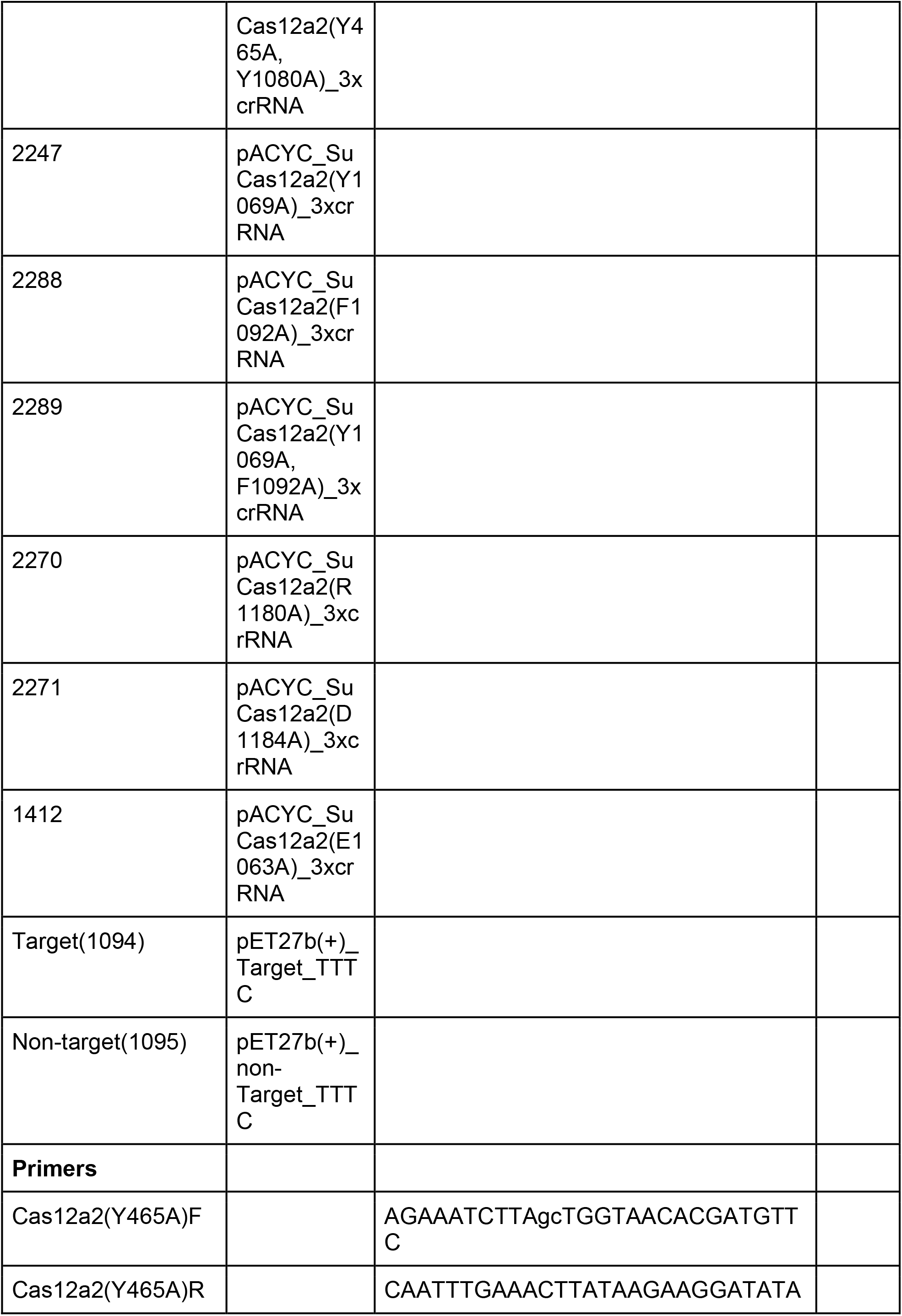
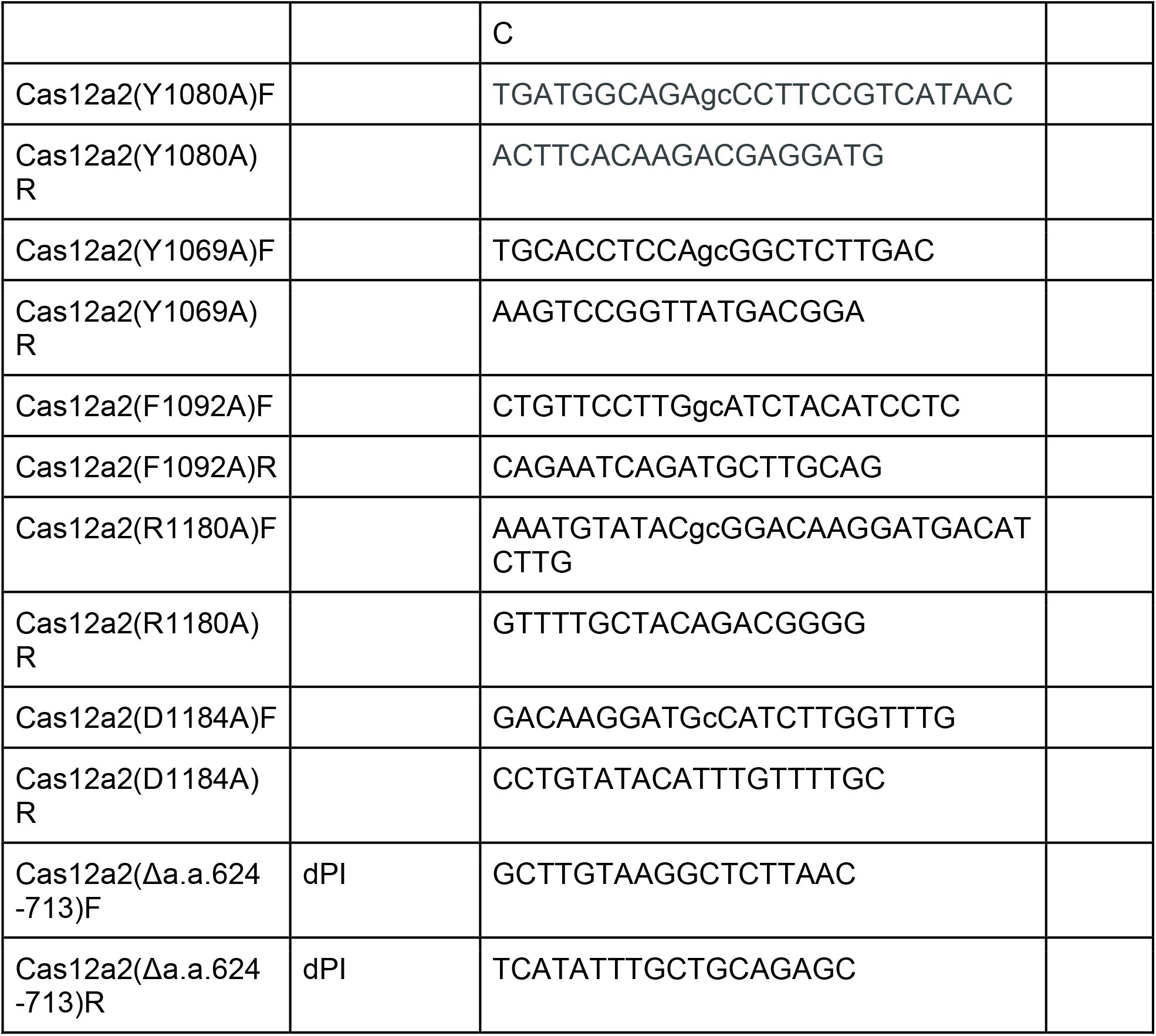
Plasmids and nucleic acid substrates.

